# The occurrence of tarsal injuries in male mice of C57BL/6N substrains in multiple international mouse facilities

**DOI:** 10.1101/2020.02.25.964254

**Authors:** Eleanor Herbert, Michelle Stewart, Marie Hutchison, Ann M. Flenniken, Dawei Qu, Lauryl M. J. Nutter, Colin McKerlie, Liane Hobson, Brenda Kick, Bonnie Lyons, Jean-Paul Wiegand, Rosalinda Doty, Juan Antonio Aguilar-Pimentel, Martin Hrabe de Angelis, Mary Dickinson, John Seavitt, Jacqueline K. White, Cheryl L Scudamore, Sara Wells

## Abstract

Dislocation in hindlimb tarsals are being observed at a low, but persistent frequency in adult male mice from C57BL/6N substrains. Clinical signs included a sudden onset of mild to severe unilateral or bilateral tarsal abduction, swelling, abnormal hindlimb morphology and lameness. Contraction of digits and gait abnormalities were noted in multiple cases. Radiographical and histological examination revealed caudal dislocation of the calcaneus and partial dislocation of the calcaneoquartal (calcaneous-tarsal bone IV) joint. The detection, frequency, and cause of this pathology in five large mouse production and phenotyping centres (MRC Harwell, UK; The Jackson Laboratory, USA; The Centre for Phenogenomics, Canada; German Mouse Clinic, Germany; Baylor College of Medicine, USA) are discussed.

## Introduction

Inbred strains of laboratory mice are used to standardise the genetic background of mutant mouse strains to reduce data variability. Produced by >20 consecutive generations of sibling mating, the controlled homogeneity of inbred strains such as C57BL/6N is accompanied by the fixing of spontaneous mutations in inbred genomes. Many monogenic mutations have been identified in inbred mouse strains, including those causing retinal degeneration in C3H strains (Schmidt, Lolley, & Racz, 1973) and age-related deafness in C57BL/6 strains (Johnson, Erway, Cook, Willott, & Zheng, 1997). The characterisation of these mutations has allowed their impact on individual research programs to be assessed and alternative genetic backgrounds used if they interfered with the primary purpose of the studies. Conversely there are reports of sporadic, low-level defects in inbred lines which are likely due to oligogenic or polygenic effects and exhibit variable penetrance thus only observed or measured in a proportion of the population of an inbred colony (Sundberg, Silva, Li, Cox, & King, 2004). These include complex behaviours such as aggression (Miczek, Maxson, Fish, & Faccidomo, 2001), hyperactivity (Võikar, Kõks, Vasar, & Rauvala, 2001), morphological anomalies such as sternal segment dislocation (Adissu, Medhanie, Morikawa, & White, 2015) and developmental defects such as hydrocephalus (https://www.jax.org/news-and-insights/2003/july/hydrocephalus-in-laboratory-mice).

Knowledge of the predisposition of mouse strains to such issues is not only essential for the care and welfare of mice but is an important consideration in phenotyping programs. It is crucial to distinguish incidental effects caused by genetic background, from outcomes arising because of an experimental paradigm (e.g. genetic mutation or physiological challenge) or a combination of background and paradigm together. The International Mouse Phenotyping Consortium (IMPC) (www.mousephenotype.org) is generating a genetically altered (GA) mouse strain carrying a null allele for each protein-coding gene in the mouse to study mammalian gene function (Brown & Moore, 2013). GA strains for this programme are generated on the C57BL/6N genetic background and phenotyping is performed at an early adult time point (up to 17 weeks) and a late adult time point (after 12-18 months) for a subset of strains. Phenotyping and husbandry protocols include the regular assessment of welfare and fitness during handling and cage-changing, and motor function during phenotyping tests (Rogers et al., 2001).

In this study, we report the recurrent observation of abnormal hindlimb morphology, accompanied by lameness, in group -housed male mice of C57BL/6N substrains. This report describes the nature of the injury and discusses possible aetiologies. We also provide an estimate of the frequency of occurrence from five large mouse genetics centres in four different countries across two continents and highlight potential consequences for projects where prolonged co-housing of male C57BL/6N mice is a necessity.

## Materials and Methods

### Ethics Statement

#### Mice were examined for tarsal injury at five mouse phenotyping centres

##### MRC Harwell

Animal studies are performed in compliance with guidelines issued by the Medical Research Council (MRC) (UK) in “Responsibility in the Use of Animals for Medical Research” (July 1993). The care and use of all mice in this study were in accordance with UK Home Office regulations, The Animals (Scientific Procedures) Act 1986 Amendment Regulations 2012 (SI 4 2012/3039), and approved by the MRC Harwell Institute Animal Welfare and Ethical Review Body.

##### The Centre for Phenogenomics (TCP)

All experimental procedures were approved by the TCP Animal Care Committee (AUP 0279) and were conducted in accordance with the guidelines of the Canadian Council on Animal Care.

##### TheJackson Laboratory (JAX)

All experimental procedures were carried out under Protocols 14004 and 11005 approved by the JAX Institutional Animal Care and Use Committee (IACUC) with NIH Office of Laboratory Animal Welfare (OLAW) assurance number D16-00170 and Accreditation AAALAC #000096.

##### German Mouse Clinic (GMC)

All animal experiments were carried out in accordance with German legal guidelines and following the approval of the responsible animal welfare authorities and the Ethics Board of the District Government of Upper Bavaria, Germany (approval number 46-2016).

##### Baylor College of Medicine (BCM)

Animal experiments were carried out in accordance with research protocol AN-5896 and approved by the BCM Institutional Animal Care and Use Committee. The Animal Welfare Assurance at BCM is approved by the Office of Laboratory Animal Welfare (OLAW), and meet the requirements of the Public Health Service Policy on Humane Care and Use of Laboratory Animals (assurance number D16-00475).

### C57BL/6N substrains used in this study

Mice used at MRC Harwell were C57BL/6NTac mice purchased from originally from Taconic Biosciences, USA and subsequently bred at MRC Harwell. Mice examined at The Jackson Laboratory are sourced from an in-house maintained colony of C57BL/6NJ. Mice used at The Centre for Phenogenomics are C57BL/6NCrl, purchased from Charles River Laboratories, USA and subsequently bred at TCP. Mice at the German Mouse Clinic were C57BL/6NTac purchased from Taconic Biosciences, Germany and C57BL/6NCrl purchased from Charles River Laboratories, Germany. Mice at Baylor College of Medicine were C57BL/6NJ originally purchased from Charles River Laboratories, USA and subsequently bred at this facility.

### Mouse housing conditions

Housing conditions in each institution are listed in TABLE S1 of supplementary figures.

All mice were given food and water *ad libitum*. Adult mice were humanely sacrificed by an overdose of anaesthetic, overdose of carbon dioxide, or by cervical dislocation (according to relevant national and local protocols and guidelines.

### Clinical examination

Routine animal care and welfare checks in all facilities involved visually inspecting the mice as part of a daily check and regular handling (typically no less than once every 14 days) during cage changing, or during phenotyping. Mice with an abnormal gait and/or locomotor deficit accompanied by abnormal hindlimb morphology and swelling or reddening of the tarsus were euthanised or selected increased observation to ensure no further deterioration in the welfare of the mouse.

### Radiography

Lateral views of the affected and contralateral tarsi were taken of representative animals under isoflurane anaesthesia by digital radiography at 26□kV for 3 □s using a Faxitron MX‐20 digital X‐ray system or a Faxitron X-Ray Model Ultrafocus 100 (both from Faxitron X‐ray Corporation, Lincolnshire, IL, USA). X‐ray images were processed using the DicomWorks software (http://www.dicomworks.com/).

### Histopathology

Immediately following euthanasia, tissues from selected mice were fixed in 10% neutral buffered formalin for a minimum of 24 hours. Following fixation, the hindlimbs (affected and contralateral) were stripped of soft tissues, decalcified in formic acid for 96 hours, and processed routinely for histopathologic evaluation. Subsequently, 4–5 μm thick, mid-sagittal sections were stained with haematoxylin and eosin (H&E) for evaluation. Representative images were acquired using an Olympus BX43 microscope with a Micropix Elite 5MP camera and Cytocam software v1.6. All histologic evaluations were performed by a board-certified veterinary pathologist.

## Results

### Clinical examination

A proportion of male mice of C57BL/6N substrains, housed in social groups were observed to have clinical signs of abnormal hindlimb morphology, together with swelling or reddening of the tarsus and often an abnormal gait. Similar numbers of C57BL/6N females were assessed and no tarsus, hind paw, or gait abnormalities were observed. Gait abnormalities in males ranged from limping with a reduced amount of weight bearing on the affected limb to complete non-weight bearing. Grossly affected tarsi showed a loss of the abrupt right angle formed from the calcaneus and the calcaneal tendon, and there was variable soft tissue swelling sometimes accompanied with redness (Figure 1). Whilst the majority of affected mice had only one abnormal hind paw, 1/21 at MRC Harwell, 4/21 at TCP and 15/58 at JAX presented with bilateral tarsal abnormalities.

**Figure 1.**
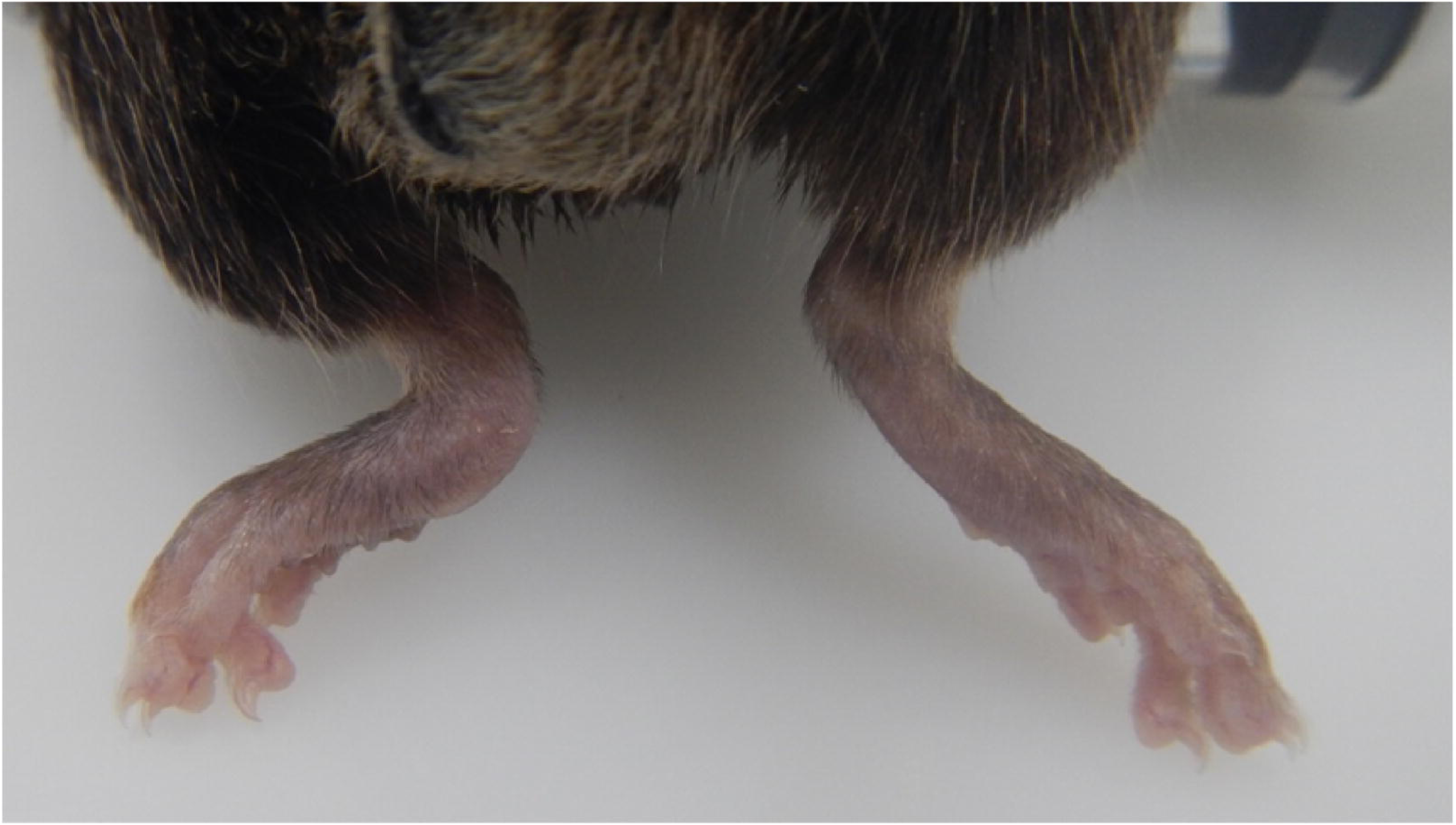
Dorsal view of unaffected right tarsus and rounded, swollen tarsus (readers left)

Radiography confirmed caudal dislocation of the calcaneus and new periosteal bone formation (Figure 2). In some animals, there was also calcification within the distal calcaneal tendon.

**Figure 2.**
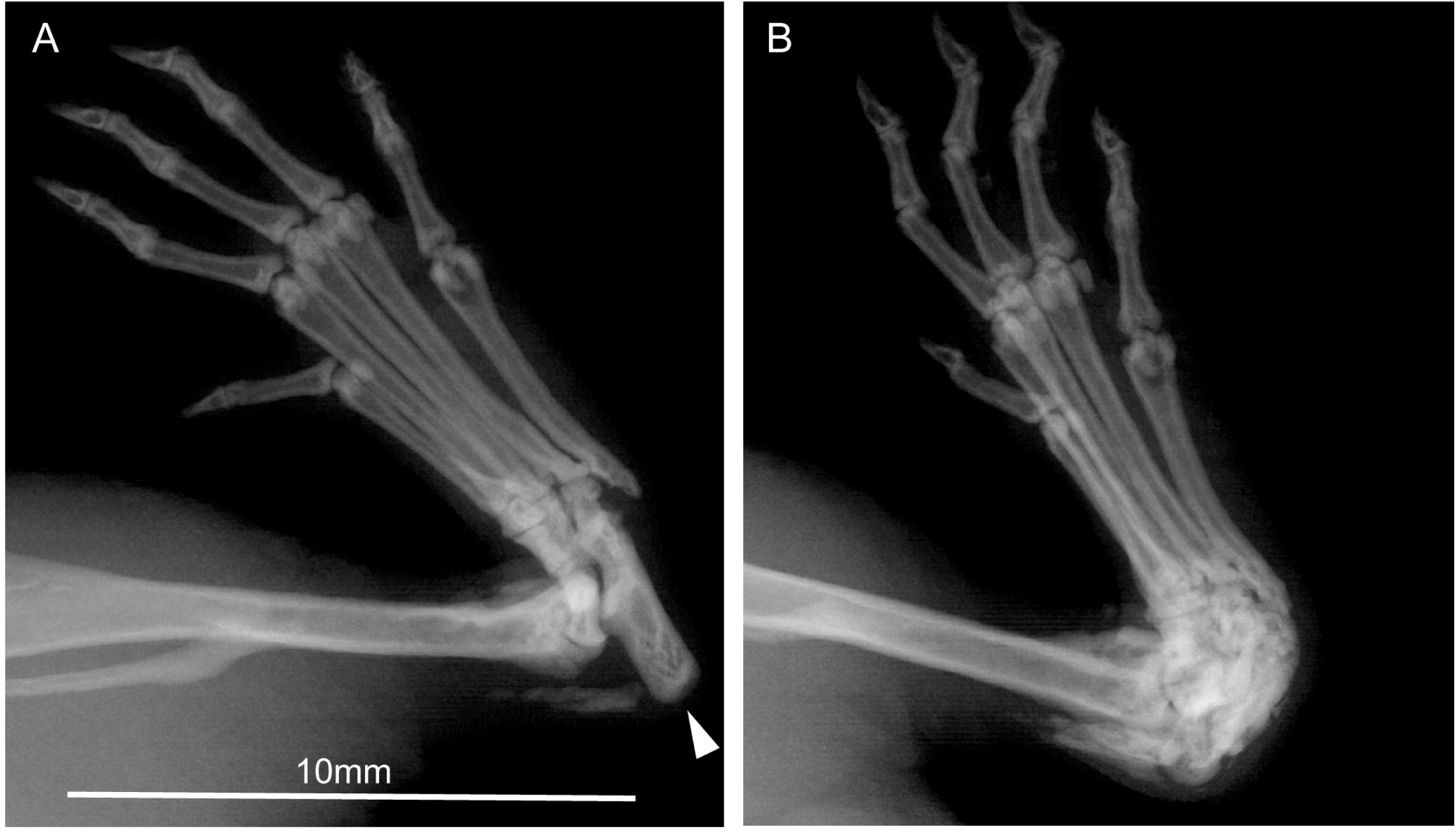
(a) Xray image of the normal position of the calcaneus (arrow) within the tarsal joint and (b) with caudo-dorsal dislocation of the calcaneus

### Histopathology

Histopathological examination identified caudo-dorsal dislocation of the calcaneus with concurrent partial dislocation and hyperextension of the calcaneoquartal joint. In more chronic lesions, the calcaneoquartal joint progressed to new bone formation (Figure 3). There was no difference in overall dislocation of the calcaneus in acute versus chronic lesions.

**Figure 3.**
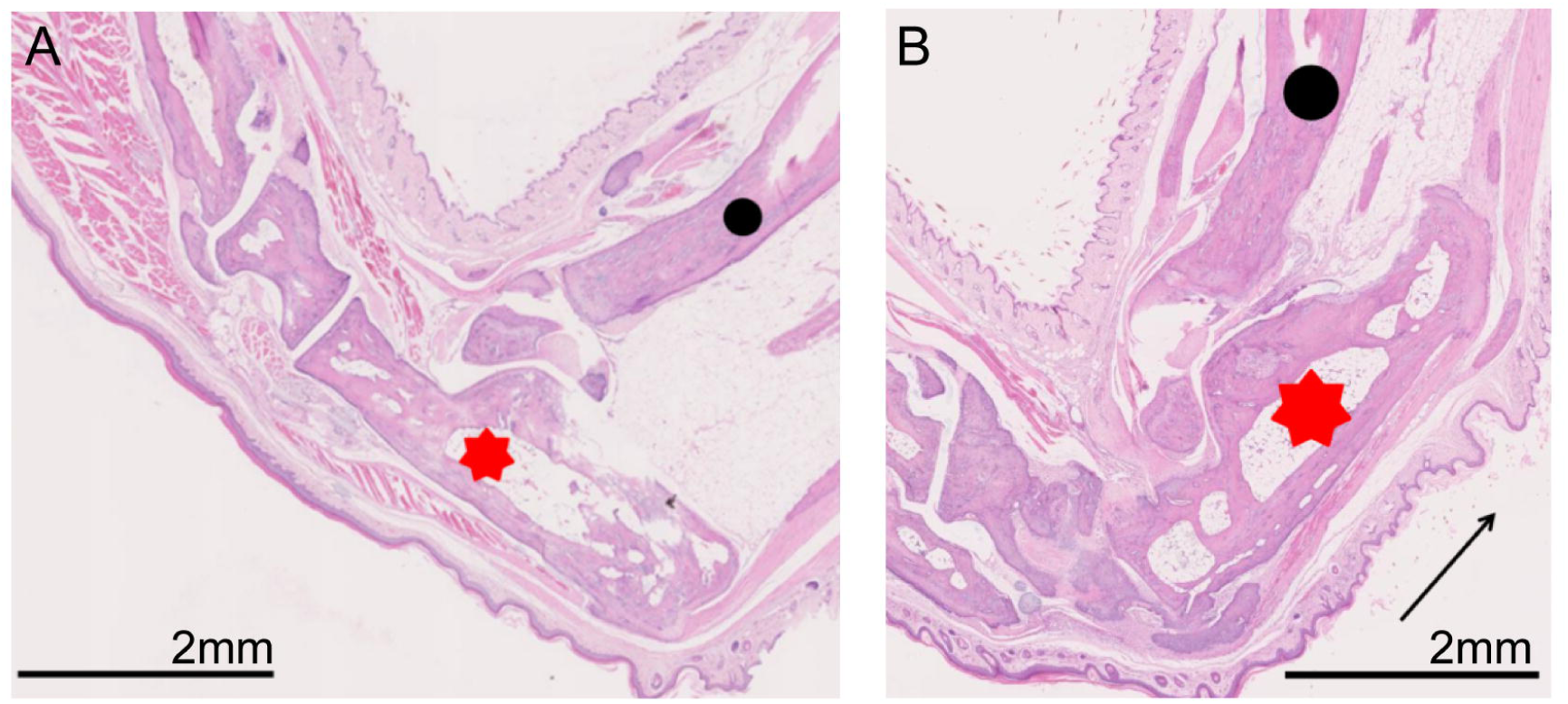
(a) The unaffected tarsus with the calcaneus (red star) forming an approximate 90° angle with the tibia (black circle), (b) Affected tarsus (red star) with dislocation of the calcaneus caudo-dorsally to form an approximate 15° angle with the tibia (black circle). The black arrow indicates the direction of movement of the calcaneus. Scale bar = 2mm.

### Frequency and variability of occurrence

C57BL/6N mice bred for the IMPC late adult phenotyping programme were examined for tarsal injuries at five international mouse research centres. These mice included GA strains from a wide range of mutant lines examined by the IMPC as well as baseline wild type controls. Due to differences in individual institutional protocol, the ages of the cohorts vary. The frequency of occurrence was between 1.7% and 12.1% of male mice examined between the ages indicated.

|  | Substrain | Number of mice (male) | Age range (weeks) | Number affected | Earliest age affected (weeks) | Frequency (%) |
| --- | --- | --- | --- | --- | --- | --- |
| The Centre for Phenogenomics , Canada | C57BL/6NCrl | 235 | 5-59 | 21 | 20 | 8.9 |
| The Jackson Laboratory, USA | C57BL/6NJ | 1440 | 4-78 | 58 | 11 | 4.0 |
| MRC Harwell Institute, UK | C57BL/6NTac | 174 | 16-59 | 21 | 18 | 12.1 |
| GMC Helmholtz Zentrum, | C57BL/6NTac and | 413 | 4-62 | 7 | 45 | 1.7 |
| Germany | C57BL/6NCrl |  |  |  |  |  |
| Baylor College<br>of Medicine,<br>USA | C57BL/6NJ | 250 | 16-52 | 30 | 20 | 12 |

### Husbandry, housing, and strains affected

Further observations were made and recorded which informed the aetiology of the incidence of the tarsal injury in C57BL/6N mice.

- Female mice: Equivalent numbers of females of the same strain which were part of the same programme of work were also examined but no similar injury was reported.
- Males in mating cages or singly-housed: No injury was observed in 584 C57BL/6Ntac males in either mating cages (with one or two females) or single-housed examined between the ages of 16 and 64 weeks (average age 24 weeks) at Harwell.

## Discussion

Here we report the observation of tarsal injury in male mice with a C57BL/6N genetic background occurring at five large and geographically-dispersed mouse facilities. These injuries were observed in a number of different mutant strains and several wild-type substrains indicating a predisposition for such lesions in mice of C57BL/6N ancestry. A similar deformity has been reported in STR/ort mice with a known genetic abnormality predisposing them to chronic arthropathy used as a model for studying osteoarthritis (Mason et al., 2001). In the STR/ort mice, lameness and hind paw deformity also affected predominantly male mice although the incidence rate was far higher and occurred from a younger age compared with our observations. The radiographic findings and histopathology are consistent with an injury caused by frequent high load tension from the calcaneal tendon through its insertion to the calcaneous leading to a breakdown of the plantar ligaments supporting the calcaneoquartal joint which are weakened in this model by a known collagen abnormality (Staines et al., 2016). There is no known underlying abnormality in the C57BL/6N strains reported here and so it is hypothesised that the lesion is caused by application of an abnormally high load/force through the calcaneal tendon because of behavioural or husbandry practices. It should be noted that all animals in this study are fed on regular maintenance or breeding diets and not on high-fat or obesity inducing regimes.

The type of lesion we identified was restricted to group-housed males and its occurrence became more prevalent as the mice aged. However, this may represent an increased opportunity for this injury to occur over time, rather than an increased predisposition/weakness in older males. The absence of any such injury in female mice socially-housed for the same experimental purposes and for the same length of time indicates that this is a sexually dimorphic effect.

As these injuries were observed in three different C57BL/6N substrains it is possible that this genetic background is predisposed to tarsal injuries. Male C57BL/6 mice are widely reported to display aggressive behaviours towards cage-mates (Lidster, Owen, Browne, & Prescott, 2019). Both threat (thrust and mounting) and aggressive behaviours (boxing, parrying, fighting) are associated with establishing and maintaining dominance hierarchies in group house male mice. Each of these behaviours involve rearing that requires repeated plantar flexion of the hind paw at the tarsus, initiated by high load tension from the common calcaneal tendon. Sporadic and frequent bouts of fighting have also been associated with an increased mechanical load on male tibiae in C57BL/6J mice (Meakin et al., 2013), a strain related to C57BL/6N. However, it is unclear whether the causal feature of the injury we observed is an inherent weakness in the tarsal joint, a consequence of a behavioural characteristic of C57BL/6N male mice, their interaction with the environment, or combinations of these factors. Significant differences in the frequency of observation of tarsal injury between centres may present any number of variances between housing and animal care regimes between the facilities. Investigations into different husbandry protocols may provide insight into ways to reduce occurrence in the future. Euthanasia following discovery of the injury described may have substantial consequences to the study being undertaken. Disruption of an established cage-group by removing and individual may lead to further perturbations in both the behaviour of the existing animals or to the experimental design itself with a reduction in data collected and the potential loss of statistical power.

In summary, this report provided a description of an injury to group-housed male C57BL/6N mice observed in five different mouse centres from studies involving large numbers of animals. It is likely that a similar incidence may occur undetected, or be being attributed to experimental protocols, in other facilities using these substrains. The implications of these findings will be study-dependent but have the potential to affect phenotyping results or cause an increase in attrition for ageing studies, resulting in insufficient animals completing the studies. The information that reported here should be used to assist future experimental design for longitudinal studies especially those involving measurements of gait and motor skills.

## Acknowledgements

We thank staff at the five facilities participating in this study for animal husbandry support, necropsy, and histology services. Dr Dona Reddiar for assistance with the manuscript.

Funding for the IMPC studies at each centre is provided by NIH grant UM1HG006348-07S2 (MRC Harwell and Baylor College of Medicine), MRC grant A410 (MRC Harwell), NIH grant UM1 OD023221 (TCP) and NIH UM1 OD0232221OD023222 (JAX), BMBF Infrafrontier grant 01KX1012 (GMC) and EU Horizon2020: IPAD-MD funding 653961 (GMC).

## Supplementary Table

**(see excel)**

## Figure legends

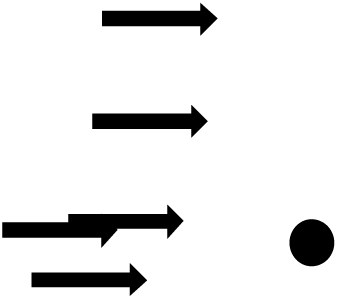

## References

Adissu, H. A., Medhanie, G. A., Morikawa, L., & White, J. K. (2015). Right Ventricular Epicardial Fibrosis in Mice With Sternal Segment Dislocation, 52(5), 967–976. https://doi.org/10.1177/0300985814552108

Johnson, K. R., Erway, L. C., Cook, S. A., Willott, J. F., & Zheng, Q. Y. (1997). A major gene affecting age-related hearing loss in C57BL/6J mice. Hearing Research, 114(1–2), 83–92. https://doi.org/10.1016/S0378-5955(97)00155-X

Lidster, K., Owen, K., Browne, W. J., & Prescott, M. J. (2019). Cage aggression in group-housed laboratory male mice: an international data crowdsourcing project. Scientific Reports, 9(1), 1–12. https://doi.org/10.1038/s41598-019-51674-z

Meakin, L. B., Sugiyama, T., Galea, G. L., Browne, W. J., Lanyon, L. E., & Price, J. S. (2013). Male mice housed in groups engage in frequent fighting and show a lower response to additional bone loading than females or individually housed males that do not fight. Bone, 54(1), 113–117. https://doi.org/10.1016/j.bone.2013.01.029

Miczek, K. A., Maxson, S. C., Fish, E. W., & Faccidomo, S. (2001). Aggressive behavioral phenotypes in mice. Behavioural Brain Research, 125(1–2), 167–181. https://doi.org/10.1016/S0166-4328(01)00298-4

Schmidt, S. Y., Lolley, R. N., & Racz, E. (1973). Cyclic-nucleotide phosphodiesterase an early defect in inherited retinal degeneration of C3H mice. Journal of Cell Biology, 57(1), 117–123. https://doi.org/10.1083/jcb.57.1.117

Sundberg, J. P., Silva, A. K. A., Li, A. R., Cox, A. G. A., & King, L. E. (2004). Adult-Onset Alopecia Areata Is a Complex Polygenic Trait in the C3H / HeJ Mouse Model. Journal of Investigative Dermatology, 123(2), 294–297. https://doi.org/10.1111/j.0022-202X.2004.23222.x

Võikar, V., Kõks, S., Vasar, E., & Rauvala, H. (2001). Strain and gender differences in the behavior of mouse lines commonly used in transgenic studies. Physiology and Behavior, 72(1–2), 271–281. https://doi.org/10.1016/S0031-9384(00)00405-4

